# Sub-Diffusive Dynamics Lead to Depleted Particle Densities Near Cellular Borders

**DOI:** 10.1101/458224

**Authors:** William R. Holmes

## Abstract

It has long been known that the complex cellular environment leads to anomalous motion of intracellular particles. At a gross level, this is characterized by mean squared displacements that deviate from the standard linear profile. Statistical analysis of particle trajectories has helped further elucidate how different characteristics of the cellular environment can introduce different types of anomalousness. A significant majority of this literature has however focused on characterizing the properties of trajectories that do not interact with cell borders (e.g. cell membrane or nucleus). Numerous biological processes ranging from protein activation to exocytosis however require particles to be near a membrane. This study investigates the consequences of a canonical type of sub-diffusive motion, Fractional Brownian Motion (FBM), and its physical analogue Generalized Langevin Equation (GLE) Dynamics, on the spatial localization of particles near reflecting boundaries. Results show that this type of sub-diffusive motion leads to the formation of significant zones of depleted particle density near boundaries, and that this effect is independent of the specific model details encoding those dynamics. Rather these depletion layers are a natural and robust consequence of the anti-correlated nature of motion increments that is at the core of FBM / GLE dynamics. If such depletion zones are present, it would be of profound importance given the wide array of signaling and transport processes that occur near membranes. If not, that would suggest our understanding of this type of anomalous motion may be flawed. Either way, this result points to the need to further investigate the consequences of anomalous particle motions near cell borders from both theoretical and experimental perspectives.

## 1. Introduction

Molecular diffusion is a fundamental process impacting almost every area of cell biology, from transport to gene regulation (as well as fields ranging from superconductor physics to finance [1, 2, 3]). But the cellular environment (cytoplasm, membrane, etc.) is a complicated and crowded place. A consequence of these environmental complexities is that proteins, mRNA, vesicles, and other diffusing entities exhibit a range of different exotic and anomalous types of random motion (see [4] for an extensive review). But how do these interactions between particles and their environment influence spatial densities? This article discusses a surprising and potentially important consequence of one particular form of sub-diffusion, Fractional Brownian Motion (FBM) and its physical analogue Generalized Langevin Equation (GLE) Dynamics, on the spatial distribution of particles near cellular boundaries.

Normal diffusion, one of the most basic nonequilibrium phenomena in nature, is well characterized by a linear mean squared displacement relationship 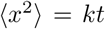 (where *k* depends on the particle and environment). It is well know however that in the complex, crowded, colloidal environment of the cytoplasm (or the cell membrane), motions of particles instead exhibit sub-diffusion^1^ where 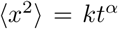 with *α* < 1. This raises the well studied questions, I) what features of the cellular environment give rise to this anomalousness, II) how can it be accounted for in stochastic models of particle motions, and III) what are its consequences. Here I focus on the latter question and consider the consequences of generalized Langevin dynamics in cellular domains.

While macro-molecular crowding has been proposed as one possible mechanism of anomalous motion [5, 6], alternative evidence has suggested simple crowding effects would reduce the speed of normal diffusion rather than introduce anomalousness [7]. Instead, some additional environmental factor is likely responsible. From a theoretical perspective there are a number of possible sources of sub-diffusion. The theory of Continuous Time Random Walks (CTRW) [8], which is typically associated with the transient caging or trapping of particles (by microtubules for example) in the filamentous cellular environment, hypothesizes that random steps are broadly distributed in time. Alternatively Fractional Brownian Motion (FBM) [9], which is usually associated with the crowded or viscoelastic nature of the cellular environment, assumes the incremental steps taken by a particle are negatively correlated due to the viscoelastic environment. Diffusion on fractals [10] also exhibit sub-diffusive characteristics, which could be particularly relevant to transport on actin or microtubule polymer networks.

Fortunately these types of motion can be distinguished from each other statistically. Motions of chromosome loci were found to obey a FBM type dynamic as opposed to CTRW [11]. Similarly, motions of tracer particles in an *in vitro* dextran solution were found to exhibit a FBM type dynamic [12]. Motions of tracer particles in an Actin network on the other hand exhibit CTRW characteristics [13]. While these types of motions are distinct and distinguishable, they are not mutually exclusive of each other [14, 15]. Insulin granules for example were found to exhibit characteristics of both [16]. This article will focus on the impact of FBM type motion on spatial densities near cellular boundaries (membrane, nucleus, etc.).

While there is a vast literature on this topic, the majority of it has sought to infer the source of anomalous motions by analyzing the statistical properties (time averaged mean squared displacement, p-variation [17, 18], turning angle distributions for example, or mean first passage time [10, 19]) of particle paths. Furthermore, most experimental studies have been limited to observing particles distant from cellular boundaries, and most theoretical studies have followed suit. Some studies [20, 21, 22] have investigated the influence of confinement on anomalous motions. They have again however focused primarily on the statistical analysis of particle paths to investigate, for example, the effects of confinement on ergodicity. A more recent investigation [23] has shown confinement of particles undergoing FBM can have significant consequences on their spatial distribution. This study however focused on transient rather than steady state dynamics and studied those dynamics on an unbounded domain (half line).

This article investigates the influence of anomalous motions on steady state particle distributions in bounded, confined domains (e.g. cells). Results show that 1) FBM / GLE dynamics lead to a significant depletion of particles near cellular boundaries, 2) these depletion effects are likely significant enough in both magnitude and spatial extent to be observed experimentally, and 3) the inherent memory dependence (i.e. increment correlations) of particle motions that are intrinsic to the basic physical hypothesis of this type of motion is responsible for this effect. This raises an important question that could be readily investigated with current super-resolution microscopy techniques: do these depletion zones exist near borders in either cellular or *in vitro* systems? If so, this would be of profound importance given the wide range of biological processes that rely on interactions with a boundary (trafficking or binding for example). If not, it may point to a significant flaw in our understanding of this form of sub-diffusive motion, both in cellular environments and more generally.

## 2. RESULTS

The effect of FBM type dynamics on the spatial localization of particles near confining borders is investigated here. The goal however is not to investigate FBM itself but rather the underlying physical assumptions that are typically associated with it. At the most basic level, FBM encodes anti-persistent motions resulting from anti-correlated increments of motion. This inherently introduces a type of history dependence into motion. While FBM accurately captures [24, 25, 26] many aspects of particle dynamics, it is not inherently a physics-based model. More recently, the Generalized Langevin Equation (GLE) [27], which is a well established generalization of the standard Langevin equation, has been proposed as a more physically grounded model with similar properties to FBM.

Both FBM and GLE dynamics will thus be investigated using simulations. Additionally, a third toy model of anti-correlated, memory dependent particle motions will be considered. This third model is considered primarily to strip out many of the complexities of the FBM and GLE models, while retaining the inherent assumption of anticorrelated increments central to these models. This approach of considering multiple, distinct models (along with multiple numerical methods) encoding similar underlying dynamics will help elucidate, in a model independent way, how the presence of anti-persistent particle motions, and the inherent memory dependence that comes along with it, influence particle densities near confining boundaries.

### 2.1. Models

A brief overview of the models used is given here. For additional details about the models themselves or the numerical implementation details, see the relevant Methods section.

#### 2.1.1. Fractional Brownian Motion (FBM)

Fractional Brownian Motion (FBM) is a generalization of Brownian motion where step increments are not independent. Let *B^H^*(*t*) denote the FBM with mean squared displacement characterized (in one dimension) by 2*Dt^2H^* where *H* is the Hurst exponent and *D* is the generalized diffusion coefficient (*H* < 1/2 corresponds to sub-diffusion). The FBM process can be generated by a fractional Gaussian noise process (FGN), given by

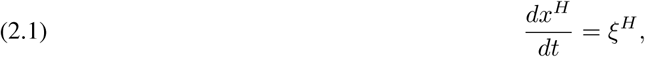

where *x^H^*(*t*) denotes the trajectory of a particle and ξ*^H^* are correlated increments with 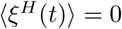 and covariance

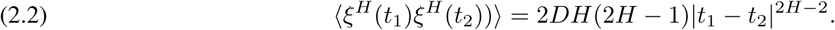

Thus FBM is a statistical model that, for *H* < 1/2, encodes the negative correlation between increments of a particle’s motion. It is this negative correlation that gives rise to anti-persistent, sub-diffusive dynamics. This model of particle dynamics will be simulated using Matlab’s wfbm package (part of the wavelet toolbox) for generating FBM. In a confined domain, it will be supplemented with a standard reflecting boundary condition.

#### 2.1.2. Generalized Langevin Equation (GLE) Motion

The GLE is a generalization of the canonical Langevin equation that can be derived from basic physical considerations of how a particle interacts with a heat bath [28]. It takes the form

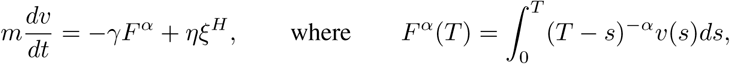

and the fluctuation-dissipation theorem [29] dictates that

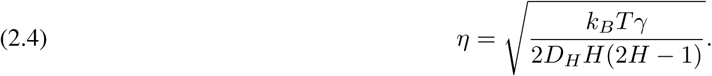

Here *F^α^* denotes a history dependent, generalized drag term, ξ*^H^* is the FGN defined above, *η* is the magnitude of that noise, and *γ* is the generalized friction coefficient. In this case, the asymptotic dynamics obey 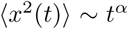 provided the Hurst exponent is *H* = 1 – α/2. The focus going forward will be on the the over-damped limit where inertia is negligible (*m* = 0). Simulations of this model for the range *γ* = 10^−10^ – 10^−9^ will be performed. This yields mean squared displacements at 60 seconds in the range 4-40 *μm*^2^. Smaller values become too computationally cumbersome due to the smaller required time steps. For further discussion of the properties of the GLE see [26].

For completeness, three methods of simulating this model will be used. The first is a lattice-free, Langevin type method that tracks the microscopic motions of particles. This method (detailed in Section 4.1) discretizes the integral in *F^α^* using standard quadrature methods. Alternatively, the predictor corrector scheme in [26] will be used. These two methods use a similar approach with slightly different implementation techniques to simulate the GLE. A third method will be used that provides a qualitatively different interpretation of the GLE as a lattice random walk [30]. This method envisions the GLE dynamics as a biased random walk where at each step, the bias due to the particle’s history is introduced by subjecting it to the force *F^α^* (see Section 4.2 for implementation details). Standard energetic / Boltzmann statistics arguments can then be used to prescribe the probability of stepping in different directions under the influence of this force. As will be shown, all three methods yield similar qualitative conclusions, though they differ in their quantitative predictions of the magnitude of boundary induced effects.

#### 2.1.3. Anti-persistent lattice random walk model (toy model)

Analysis of the FBM and GLE results indicates that the history dependence inherent in the two models is vitally important to the observed results. However both models have numerous complexities and assumptions embedded in them. To assess the influence of this memory dependence on spatial localization of particles near boundaries, a simple lattice random walk model is developed (see Section 4.3) where the probability of right / left steps depend on the number of right / left steps taken over the previous *T_m_* steps. For example, if most of the walker’s steps were to the right, a leftward bias is introduced and vice versa. The parameter *T_m_* encodes how long into the past this memory process remembers and an additional bias parameter is included that incorporates the strength of the history dependent bias. This model will be used to determine the influence of memory, and the length of that memory, on spatial distributions.

### 2.2. Modeling Results

First, simulations of spatial particle densities at steady state were performed for different values of α for both FBM and the GLE. They demonstrate a surprising result: when particle motions are sufficiently anomalous, a substantial depletion zone in the steady state distribution appears near the cellular border (Figure 1 a,c). Alternative simulations of the GLE using the previously developed numerical method [26] show a similar effect that is even larger in magnitude (Figure 2a). Furthermore, the lattice Monte Carlo based GLE simulations in both 2D (Figure 3) and 1D (Figure 4b) show similar results. Finally, the simple anti-persistent lattice random walk model shows similar depletion zones (Figure 1d) that become more significant as the strength of the bias effect increases. This effect is however only prominent when the anomalous exponent (α) is well below one (or in the toy model case when the bias is sufficiently strong). In all cases, when the exponent approaches *α* =1, these depletion zones shrink and the expected uniform densities appear again. In combination, these results suggest that the more sub-diffusive the motility, the more substantial this depletion effect becomes.

**Figure 1.**
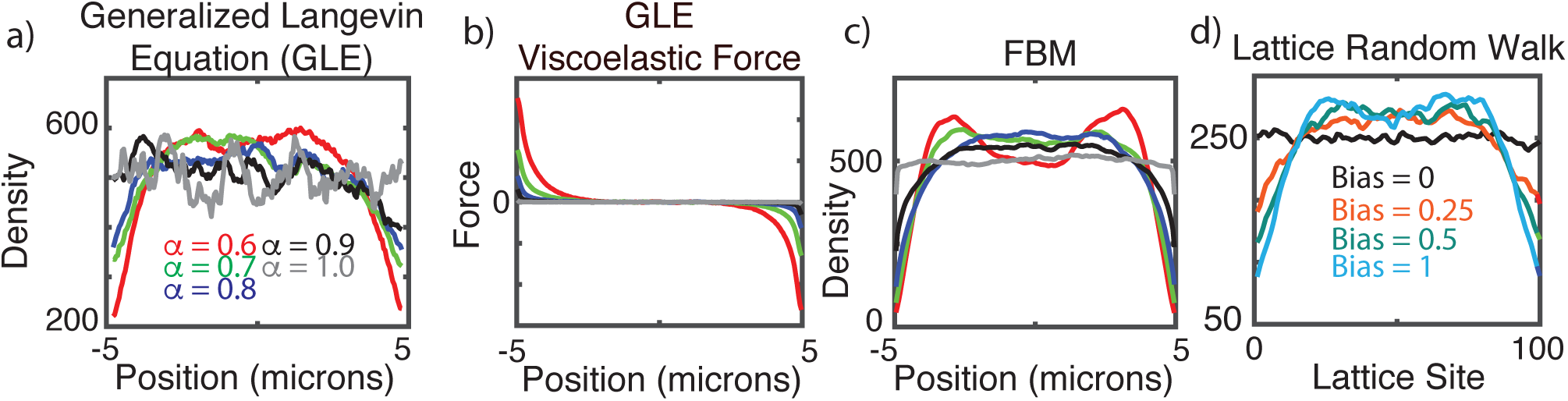
Anomalous motions lead to depletion of densities near boundaries in a model independent fashion. **(a)** Simulated density profile (in 1D) for the GLE with 5,000 particles and varying values of a at a fixed value of *γ* = 10^−9^. Densities calculated after T=250 sec with a time-step of 1/100 sec. **(b)** Spatial profile of the friction force *F^α^* for the simulations in (a). The force profile is averaged over all particles and all times in the interval *T* ∊ [240, 250]. **(c)** Simulated density profile for the FBM with 5,000 particles and varying values of α at a fixed value of *D* = 1. Densities calculated after T=6000 sec with a time-step of 1/100 sec. **(d)** Simulated density profile for the toy model with different biasing strengths. 20,000 particles were simulated on a lattice with 100 sites for 20,000 time steps.

**Figure 2.**
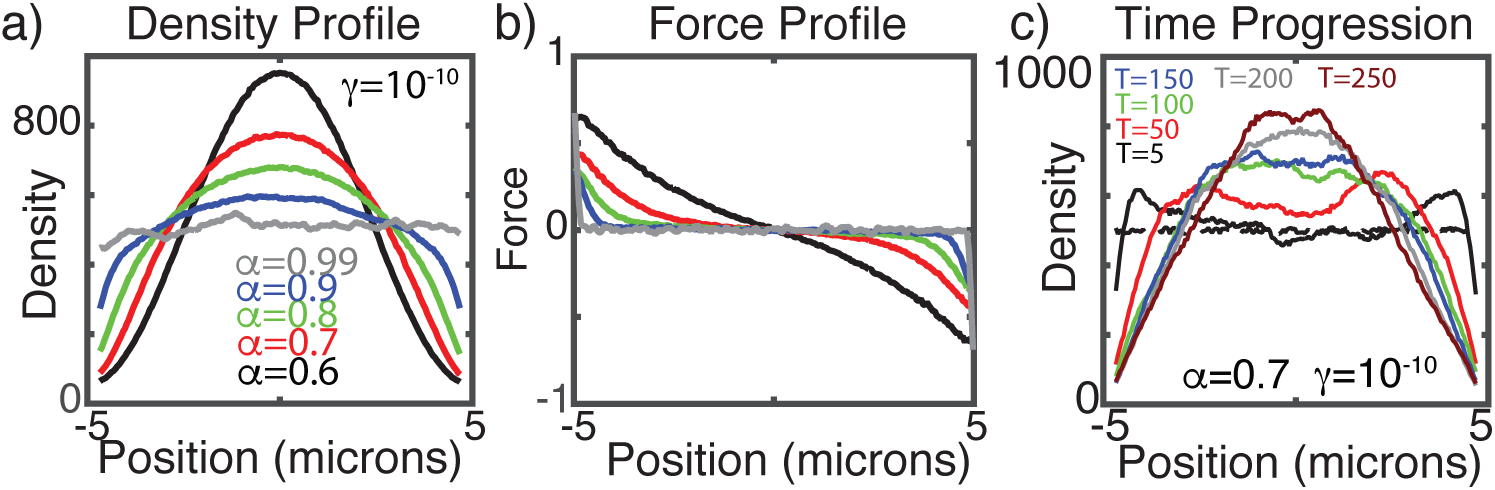
Verification of results with a different numerical scheme. **(a, b)** Simulations from Figure 1 (a,b) were repeated with the numerical method from [26]. Similar results (not shown) are observed for *γ* = 10^−9^. **c)** Simulated time progression from an initially homogeneous density state (dashed line) for the α = 0.7 case from (a).

**Figure 3.**
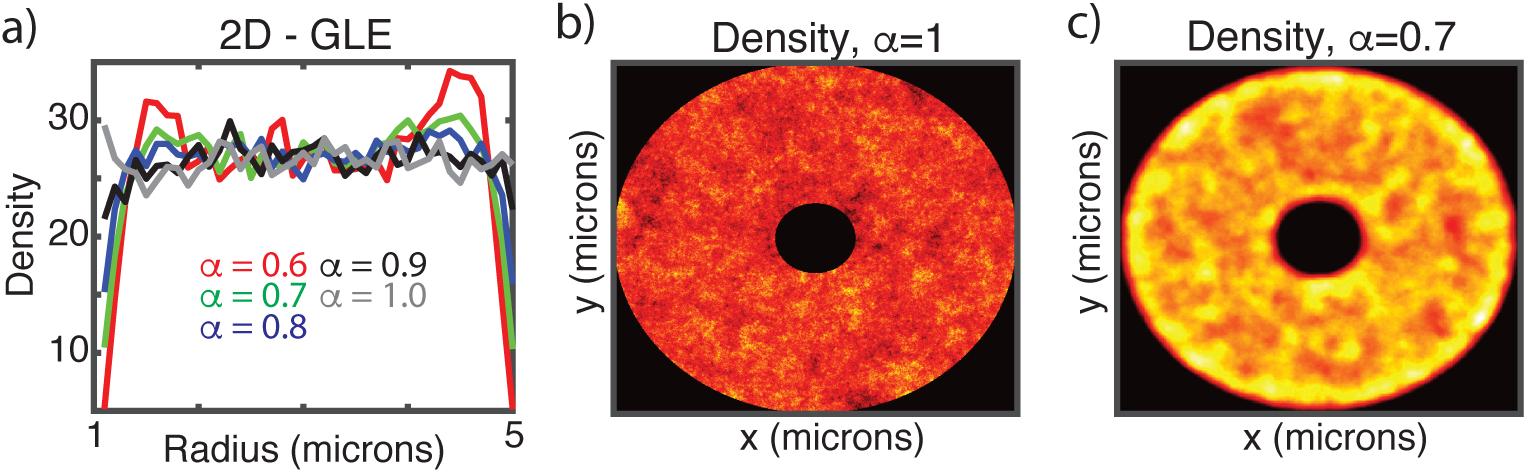
Verification of results in 2D with a lattice random walk implementation of GLE dynamics. **(a)** Simulated GLE on a 2D annulus using the lattice based method, demonstrating dependence of depletion layers on α. Plots show density as a function of radius where *r* = 1 denotes the nuclear membrane and *r* = 5 the cell membrane. Simulations were calibrated to insulin granule motility data from [33] so that the MSD after 4 minutes is ∼ 1*μm*^2^ for α = 0.7. 20,000 particles were simulated for 1000 sec with *dt* = 1/100 sec. **(b, c)** 2D spatial density map at the end of these simulations illustrating depletion near the nuclear and cell border for α = 0.7. Red / yellow indicate low / high density respectively.

**Figure 4.**
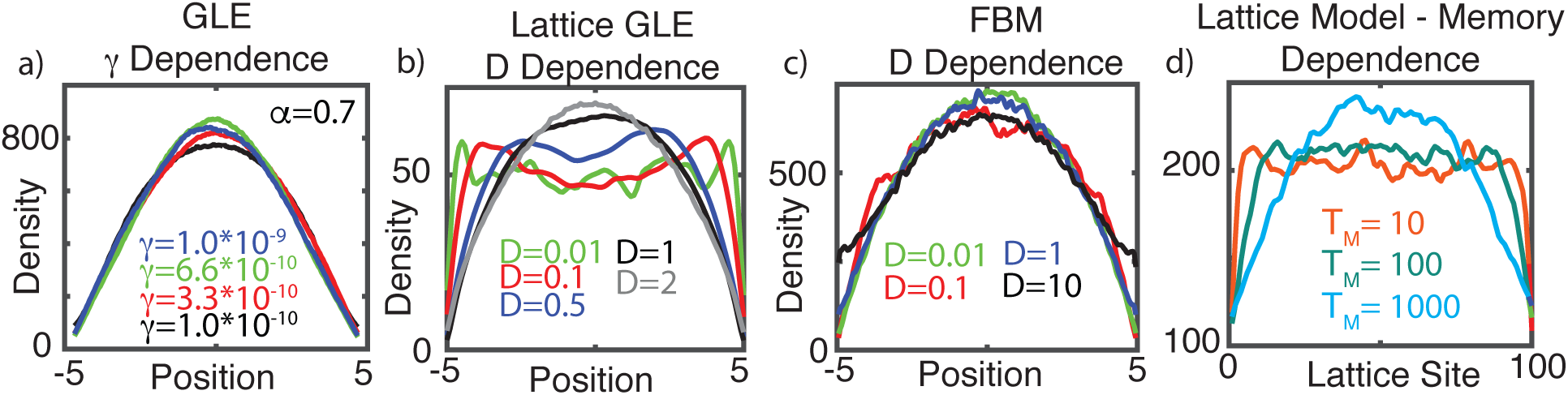
Increasing particle mobility either enhances or has little influence on boundary depletion effects. **(a)** Simulated dependence of depletion layers on particle mobility for the GLE (in 1D) using the method from [26] and simulation specifics the same as in Figure 1a. **(b)** Simulated dependence of depletion layers on particle mobility for the 1D GLE using the lattice based method. D is a proxy for the generalized diffusion coefficient in this method and these values produce MSD after 1 minute that range from 0.04 ‱ 7*μm*^2^. 5,000 particles were simulated for 500 sec with a timestep of *dt* = 1/400. **(c)** Dependence of depletion layers on mobility for 1D FBM using α = 0.7 and similar simulation details as Figure 1c. **(d)** Dependence of depletion layers for the toy lattice model on the length of the memory process (e.g. *T_m_*).

Next, the influence of particle mobility on the presence of these zones was assessed. For FBM, the spatial profile appears to be nearly unaffected by four orders of magnitude variation in *D* (Figure 4b). For the GLE, the effect appears to be insensitive to varying *γ* (Figure 4a). For the lattice version of the GLE (Figure 4b), varying *γ* has a clear effect with faster particle motion (smaller *γ*) yielding substantively more significant depletion effects. In conclusion, while there are discrepancies between results in Figures 4 a-c, all demonstrate that increasing particle mobility either increases or has no detectable effect on this boundary depletion. Thus, increasing particle mobility would not abrogate the effect as might be expected given the homogenizing tendency of diffusion.

Finally, the influence of “memory length” on these boundary depletion effects was assessed. Simulations of the anti-persistent lattice random walk model with different memory lengths (Figure 3c) show that the longer the memory, the larger the steady state depletion effect. Note that this model does not explicitly encode any spatial or temporal scale and there is no explicit α encoding the level of anomalousness. The only aspect of this model capable of producing heterogeneity is the anti-persistent memory incorporated into it.

### 2.3. Explanation of boundary depletion effects in the GLE

The presence of this depletion effect in the toy model and its dependence on memory length (*T_m_*) provides a clue as to the source of this boundary depletion. Consider a particle observed near the right domain boundary. It is impossible for such a particle to have taken a persistent path from the right of its current location due to the boundary. More generally, a particle observed near the right boundary is more likely to have taken a majority of rightward steps rather than leftward steps to get to that location. Thus a particle observed near a boundary will necessarily have a biased history. Given the anti-persistence of all of these models, this biased history would be expected to introduce a net bias in the motion of any particle near a boundary. Furthermore, the closer a particle is to the boundary, the more substantial that bias would be. Thus a particle near a boundary would feel (on average) a net bias to move away from the boundary even without any explicit boundary influence. Meanwhile, a particle well away from the boundary would not be expected to experience (on average) any net bias.

How does this explanation translate to the physics-based GLE model, which does not encode an explicit history dependent motion bias? In the GLE, the generalized friction term *F^α^* encodes the history dependence of particle’s motion. The hypothesis based on the above mechanism is that a particle near a boundary is likely to (on average) have a biased history. If true, that should manifest as a heterogeneous force profile (*F^α^*) at steady state

To test this, *F^α^* was computed and averaged over every particle over a 10 second time frame to produce a force map. Using both the simulation method developed here (Figure 1b) and the method in [26] (Figure 2b), the resulting force profile is heterogeneous (at steady state) as expected. Away from the boundaries there is no net friction force when averaged over all particles. There is however a net positive (respectively negative) force near the left (resp. right) boundary. Thus, a particle near the left boundary experiences (on average) a generalized friction force pushing it toward the right (and vice versa for the right boundary). This is consistent with the toy model results, suggesting that the simple fact that a particle observed near a boundary necessarily has a biased history will introduce these depletion effects. Furthermore, it suggests that super-diffusion, where motion increments are positively correlated, would be expected to introduce a commensurate boundary enrichment, as observed in [23] for FBM.

In conclusion, the anti-persistent “memory” of these processes leads to the formation of depleted density zones near cell boundaries. This effect is independent of the specific model encoding these dynamics or numerical method used for simulation. Furthermore, this effect is either insensitive to, or exacerbated by, increases in particle mobility. Its magnitude also increases as the memory length increases. Finally, it is not the statistical or physical details of these models that are responsible for this observation. Rather it is the anti-persistent memory, which is at the very heart of FBM / GLE type sub-diffusion, that is responsible.

## 3. Discussion

Results here indicate that Fractional Brownian Motion (FBM) / Generalized Langevin Equation (GLE) dynamics interact with confining boundaries to produce significant density depletion zones near those boundaries. This effect is significant when the sub-diffusive exponent is *α* < 0.8, which is a relevant range for the motion of numerous biomolecules (α ∼ 0.7 has been commonly observed, see [4]). Furthermore, this effect is present in multiple models that encode the basic underlying assumptions of FBM (e.g. anti-correlated increments) and appear using multiple simulation methods for the GLE.

This is of potentially profound importance. How proteins, vesicles, and other molecules localize near various boundaries in the cell is vital to numerous cellular processes. Transport of transcription factors across the nuclear membrane is vital to gene regulation [31,32]. Localization of vesicles near the cell membrane is a precursor to insulin secretion in beta cells [16, 33]. Cycling of proteins on and off of the membrane is critical to numerous regulatory processes (cell polarity or wound healing for example) [34, 35, 36, 37]. Thus it is of vital importance to understand how motions of particles in the complex cellular environment influence localization near these boundaries.

These results raise a number of important questions. Do these depletion zones exist near borders in either cellular or *in vitro* systems? If so, this effect could fundamentally alter the dynamics of any process that requires the localization of substances near such a border. Alternatively, if such effects are not present, what does that say about our understanding of this form of anomalous motion (sub-diffusive motion that is not of CTRW type)? Is the incorporation of history dependence into particle dynamics fundamentally flawed from a biophysical perspective? Or is this simply an issue of properly specifying boundary conditions? Answering these questions may have wide-ranging implications to our understanding of the micro-rheology of cellular environments [38] and how anomalous particle motility is influenced by the viscoelastic cellular environment.

Fortunately the experimental techniques necessary to address these questions are readily available. Existing superresolution microscopy should be adequate to either measure the spatial densities of tracer particles in confined domains, or alternatively look for deviations in the properties of paths of particles (motion biases for example) close to versus far from confining boundaries. Initially, *in vitro* systems such as tracer particles in dextran (e.g. [12] or similar) may be a more promising starting point due to the numerous biophysical complexities associated with cellular membranes.

Additional theoretical work is also needed. While all simulation studies shown here demonstrate this memory dependent depletion effect, different models and numerical methods indicate differing magnitudes and influence of particle mobility on that magnitude, which may impact the experimental approach needed to search for this. Thus further study of this discrepancy is an important step going forward. Given the focus on densities, a potential alternative approach to study this issue is using a continuum fractional Fokker Planck [39, 40] approach rather than stochastic simulation.

In conclusion, this article demonstrates a significant effect of reflecting boundaries (e.g. cellular membranes) on the steady state spatial distribution of sub-diffusive particles obeying FBM / GLE dynamics. This depletion effect is a theoretically robust and experimentally testable prediction that is an intrinsic feature of the underlying FBM / GLE theory rather than the mathematical implementations of that theory. The questions that remain are: how significant is this effect and more importantly, is it truly present in relevant systems?

## 4. Methods

### 4.1. Numerical simulation of the over-damped generalized Langevin equation

The purpose here is to derive a new method for simulating trajectories from the stochastic GLE. Consider the following form of the over-damped GLE

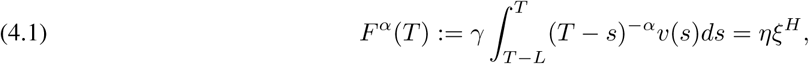

where *L* is included as a memory length for generality. Here *F^α^* denotes the generalized friction force, *α* is the exponent encoding the anomalous nature of diffusion such that *MSD* ∼ *t^α^*, *H* = 1 – α/2 is the Hurst exponent, and ξ*^H^* are the fractional Gaussian noise (FGN) increments that generate the necessary fractional Brownian motion (FBM) *B^H^*. *γ* and *η* are the generalized friction coefficient and noise amplitude respectively. *L* is the memory length, which accounts for the fact that velocities in the distant past do not contribute to damping.

To begin, let Δ*t* be the time step to be utilized and define *t_j_* = *j*Δ*t* along with *y_j_* to be the position at time *t_j_*. Discretizing the velocity in Equ. (4.1) yields

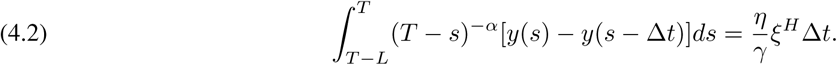

Substituting *T* = *t*_*n*+1_, the integral arising from the force kernel can now be approximated using a standard finite differencing of *y*′(*s*) as

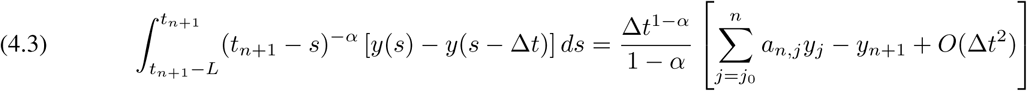

where

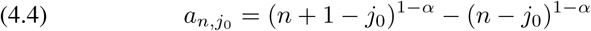

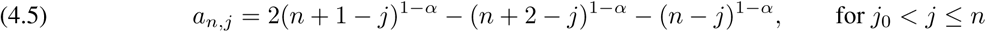

Here we have utilized that *y*(*s*) – *y*(*s* – Δ*t*) = *y*_*j*+1_ – *y_j_* + *O*(Δ*t*^2^) on the interval s ∊ [*t_j_*, *t*_*j*+1_].

It is now simple to derive an expression for the position of the particle at t_*n*+1_

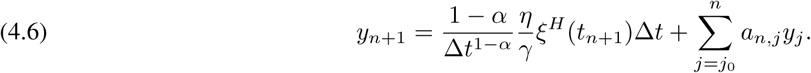

Note that the term ξ*^H^* (*t*_*n*+1_) represents a single FGN increment over time Δt. These increments were synthesized by numerically differentiating (e.g. ξ*^H^* (*t*_*n*+1_)Δt = *B^H^*(*t*_*n*+1_) – *B^H^* (*t_n_*)) a FBM path *B^H^* constructed using Matlab’s built in generator (wfbm). The Hurst exponent has been used here to denote the level of anomalousness as this is what this (and other similar) FBM generator requires. Figure 5 demonstrates that this numerical scheme produces the appropriate mean squared displacement slopes in log - log space for varying values of α. This method will be used in addition to that in [26] to ensure results are not the result of scheme specific numerical effects.

**Figure 5.**
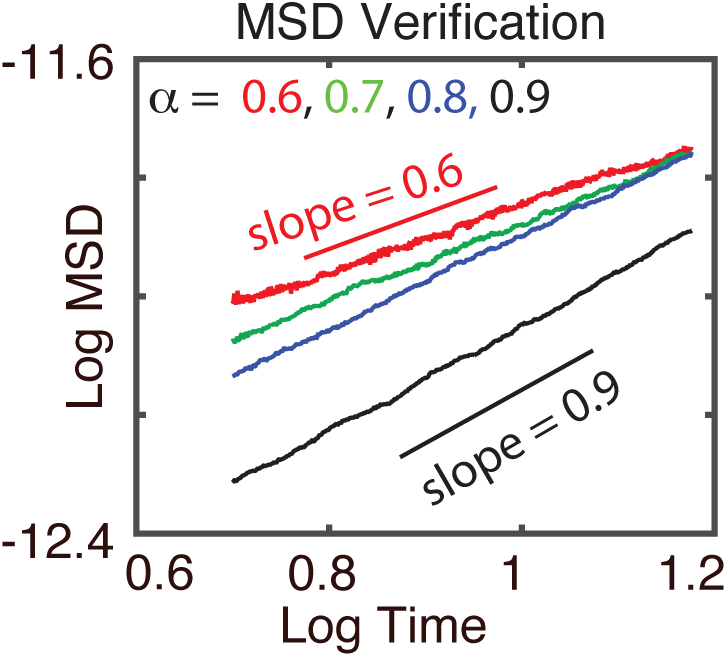
Verification of numerical simulation method. Verification that the numerical method in Section 4.1 produces the appropriate MSD scaling.

For this study, we simulate the GLE on a fixed, bounded domain *y* ∊ [–*L*, *L*] where *L* = 5 microns. In order to account for boundaries, this scheme is augmented with a standard reflecting boundary so that *y_n_* → *y_n_* – 2|*y_n_* – *sgn*(*y_n_*)*L*| if |*y_n_*| > *L* where *sgn*(*y_n_*) indicates the sign of the particle’s position.

### 4.2. Numerical simulation of the Kinetic Monte Carlo (KMC) interpretation of the GLE

Here we follow the approach of [30]. This is a lattice based approach to simulating sub-diffusive dynamics governed by the GLE. The key to this approach is to treat sub-diffusion motion as a biased lattice random walk where the bias at any point in time is determined by the particle’s trajectory history. Full details can be found in [30], but I briefly describe the approach.

Consider a one dimensional scenario where Δ*x* represents the spacing of the lattice grid points. Suppose at time *t_n_* the particle is at location *y_n_* with a past trajectory history {*y_i_*}*_i_*_=1,…,*n*_. Given this trajectory, the particle will be subjected to the viscoelastic restoring force *F^α^* (Equ. (4.1)). Define *P*_*n*,±_ to be the probabilities of taking a step to the right (+) or left (−) on this lattice. Then Boltzmann statistics dictate that

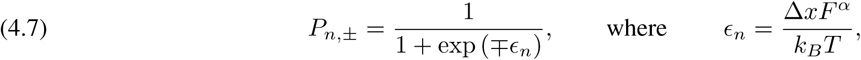

and *F^α^* is suitably evaluated over the particle’s history. In this way, the force *F^α^* is creating an effective force that the particle must fight and the *P*_*n*,±_ utilizes Boltzmann statistics and the work required to move against that force to calculate dynamically changing stepping probabilities. For full implementation details see [30].

Once again, particle trajectories are simulated on a fixed, bounded domain *y* ∊ [–*L*, *L*] where *L* = 5 microns. Standard reflecting boundary conditions are once again implemented so that any step that attempts to take a particle onto the domain’s border is aborted and the particle is replaced at its original location for that time step. This method can easily be extended to two spatial dimensions as well. At each time step, a random number is drawn to determine which spatial dimension a step will be attempted in (each dimension has equal probability). The 1D method previously described is then used to determine the direction of the step in that dimension.

### 4.3. Anti-persistent lattice random walk model (toy model) - Additional details

Here a highly simplified toy model of viscoelastic sub-diffusive motion is constructed that incorporates the characteristics of the GLE without the complexities of the fluctuation-dissipation theorem or Boltzmann statistics that are necessary for a more physically grounded model.

The most basic assumption embedded into the viscoelastic theory of sub-diffusive motion is that a particle’s environment endows it with a kind of anti-persistent memory. That is, the longer a particle moves in a particular direction, the less likely it is to continue moving in that direction in the near future. Here a toy model of a lattice random walk is constructed with this, and only this feature.

Consider a 1D lattice random walk on a lattice with sites *i* =1… *N* (N=100 for specific simulations). Define the position of the random walker to be *y_n_* and define the sign of each step to be *s_n_* = *y_n_* – *y*_*n*–1_. That is, *s_n_* = ±1 depending on whether the previous step was to the right or left. Now define the probability of taking a step in the positive direction at time *n* +1 to be

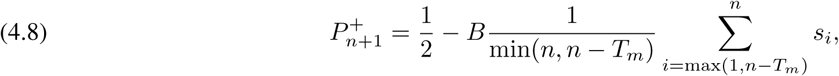

where *T_m_* denotes the memory length of the process and *B* > 0 is the strength of the memory induced bias. Note that when *B* = 0, this prescribes a pure diffusion process. The probability of stepping to the left will be 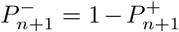. Thus if *T_m_* = 100, this summation will average over the previous 100 steps. This function averages over the previous *T_m_* steps to produce a history dependent probability of stepping to the right. Consider the three following possible particle histories in the simple case where *B* = 0.5. If all possible steps were to the right, then 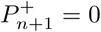. If all possible steps were to the left, then 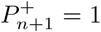. If half of all steps were to the right / left, then 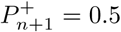. This is a simply way to incorporate the anti-persistence inherent in the GLE without its additional complications. A standard reflecting boundary condition is once again used. Any attempt to step off of the bounded lattice is aborted and the particle is replaced at its original position.

This is not intended to be a replacement model of viscoelastic sub-diffusion. Rather it is a toy model that will be used to assess the influence of anti-persistent particle memory and the length of that memory.

## Footnotes

1 In some cases, particularly where molecular motors are involved, motions can be super-diffusive with α > 1.

